# Intracortical microstimulation pulse waveform and frequency recruits distinct spatiotemporal patterns of cortical neuron and neuropil activation

**DOI:** 10.1101/2022.01.14.476351

**Authors:** Kevin C. Stieger, James R. Eles, Kip A. Ludwig, Takashi D.Y. Kozai

## Abstract

**Background:** Neural prosthetics often use intracortical microstimulation (ICMS) for sensory restoration. To restore natural and functional feedback, we must first understand how stimulation parameters influence the recruitment of neural populations. ICMS waveform asymmetry modulates the spatial activation of neurons around an electrode at 10 Hz; however, it is unclear how asymmetry may differentially modulate population activity at frequencies typically employed in the clinic (e.g. 100 Hz).

**Objective:** We hypothesized that stimulation waveform asymmetry would differentially modulate preferential activation of certain neural populations, and the differential population activity would be frequency-dependent.

**Methods:** We quantified how asymmetric stimulation waveforms delivered at 10 Hz or 100 Hz for 30s modulated spatiotemporal activity of cortical layer II/III pyramidal neurons using in vivo two-photon and mesoscale calcium imaging in anesthetized mice. Asymmetry is defined in terms of the ratio of the leading phase to the return phase of charge-balanced cathodal- and anodal-first waveforms.

**Results:** Neurons within 40-60μm of the electrode display stable stimulation-induced activity indicative of direct activation, which was independent of waveform asymmetry. The stability of 72% of activated neurons and the preferential activation of 20-90 % of neurons depended on waveform asymmetry. Additionally, this asymmetry-dependent activation of different neural populations was associated with differential progression of population activity. Specifically, neural activity increased over time for some waveforms at 10 Hz, but decreased more at 100 Hz than other waveforms.

**Conclusion:** These data demonstrate that at frequencies commonly used for sensory restoration, stimulation waveform alters the pattern of activation of different but overlapping populations of excitatory neurons. The impact of these waveform specific responses on the activation of different subtypes of neurons as well as sensory perception merits further investigation.

## Introduction

Intracortical microstimulation (ICMS) is a powerful tool to provide sensory feedback and restoration during control of neural prosthetics(1-6). Despite demonstrated clinical efficacy, the mechanisms responsible for driving neural activation during electrical stimulation are still unclear(7-11). Importantly, the stimulation-induced population activity depends on the electrical and geometrical properties of the cellular composition (excitatory/inhibitory neurons, soma, passing axons, glia)(10-18) proximal to the electrode. This complexity and relative randomness of cellular subtypes in the proximity of the electrode presents a challenge in understanding how electrical stimulation parameters result in different sensations(9, 19). For example, because neuron subtypes and neuron subnetwork connectivity contribute to specific aspects of sensory encoding(20), improving our understanding of how stimulation parameters influence activity in neural subpopulations may increase the ability to selectively elicit diverse sensations across an electrode array.

The shape and polarity of the stimulation waveform has been suggested to modulate the selectivity of activation at the soma, axons, or dendrites around the electrode(13, 14, 21-23). We previously demonstrated that the spatial activation of neurons around the electrode during 10 Hz-stimulation depends on the ratio of the leading phase relative to the return phase of biphasic waveforms, referred to as waveform asymmetry(24). However, stimulation frequency can also shape recruitment selectivity(13, 25, 26) and perception(1, 6, 27). Additionally, we demonstrated that during 100 Hz-stimulation the activity of some neurons subsides during the pulse train after a short (1-2s) period of activation, whereas activation remains stable for others(28-30). This “onset” effect is thought to be mediated by inhibitory network recruitment in part because inhibitory neurons may entrain to 100 Hz-stimulation longer than excitatory neurons(28, 29). Therefore, investigating the temporal activation of cortical neurons during 100 Hz-stimulation with different waveforms could elucidate how the asymmetry of stimulation waveform modulates recruitment patterns(13, 23, 24, 29).

In this study, we combine two-photon and mesoscale calcium imaging of pyramidal neurons (Thy1+)(31) in the mouse somatosensory cortex to test if there is an interaction between waveform asymmetry and stimulation frequency on the spatiotemporal patterns of neuronal activity. Specifically, we hypothesize that waveform asymmetry-dependent activation of neural populations also depends on frequency. Ultimately, we identify key dynamics that provide insight into the mechanisms of how ICMS waveform asymmetry recruits different but overlapping populations of neurons and highlight the need to better understand the contribution of inhibitory neurons.

## Materials and methods

### Animals and surgical preparation

All experiments were performed in 6 mature, male, (>8 weeks, >28g) Thy1-GCaMP6s mice (C57BL/6J-Tg(Thy1 GCaMP6s)GP4.3Dkim/J; Jackson Laboratories, Bar Harbor, ME)(31) in accordance with the National Institutes of Health Guide for Care and Use of Laboratory animals and were approved by the Division of Laboratory Animal Resources and Institutional Animal Care and Use Committee at the University of Pittsburgh.

Cranial window preparation for calcium imaging was carried out as previously published(32-35). Briefly, animals were anesthetized with a ketamine/xylazine mixture (75mg/kg, 7mg/kg respectively) and ketamine (40 mg/kg) was used to maintain sufficient anesthetic plane. Animals were head-fixed in a stereotaxic frame on a heating pad with continuous O_2_ supply. Two stainless-steel bone screws cemented over bilateral motor cortices acted as ground/reference electrodes, and bilateral craniotomies (3-4mm diameter) were drilled over the visual and somatosensory cortices. Sterile saline was used to maintain brain hydration. Animals received an intraperitoneal injection of SR-101 to identify insertion locations with limited vasculature and to verify limited tissue displacement.

### Two-photon and mesoscale imaging

Mesoscale imaging (MVX-10 epifluorescence microscope; Olympus, Tokyo Japan) was performed to visualize stimulation-induced activity on a large spatial scale (3.7 × 3mm)(24, 36, 37). Time series images were captured at 20 frames per second (fps) using a CCD camera (Retiga, R1 18 imaging). A two-photon microscope (Bruker, Madison, WI) equipped with an OPO laser (Insight DS+, Spectra Physics, Menlo Park, CA) at 920 nm and a 16x 0.8NA water immersion objective (Nikon, Melville, NY) was employed to answer questions related to the different temporal calcium dynamics with cellular resolution (407 × 407 μm, 0.8 μm/pixel; 30 fps).

### Electrode implantation

Electrochemical activation of iridium oxide electrode sites was carried out on acute, 16-site, single-shank Michigan style electrode arrays (A1×16-3mm-100-703; NeuroNexus, Ann Arbor Michigan) to increase the charge storage capacity and lower 1kHz impedance to <500 kOhm(38). Electrodes were then implanted into layer II/III of the somatosensory cortex under a two-photon microscope or a mesoscale microscope at a 30° angle with an oil hydraulic Microdrive (MO-81 Narishige, Japan) using previously optimized methods (Fig. 1)(24, 28, 29, 33). Sterile saline was used throughout the procedure to keep the brain hydrated.

**Figure 1:**
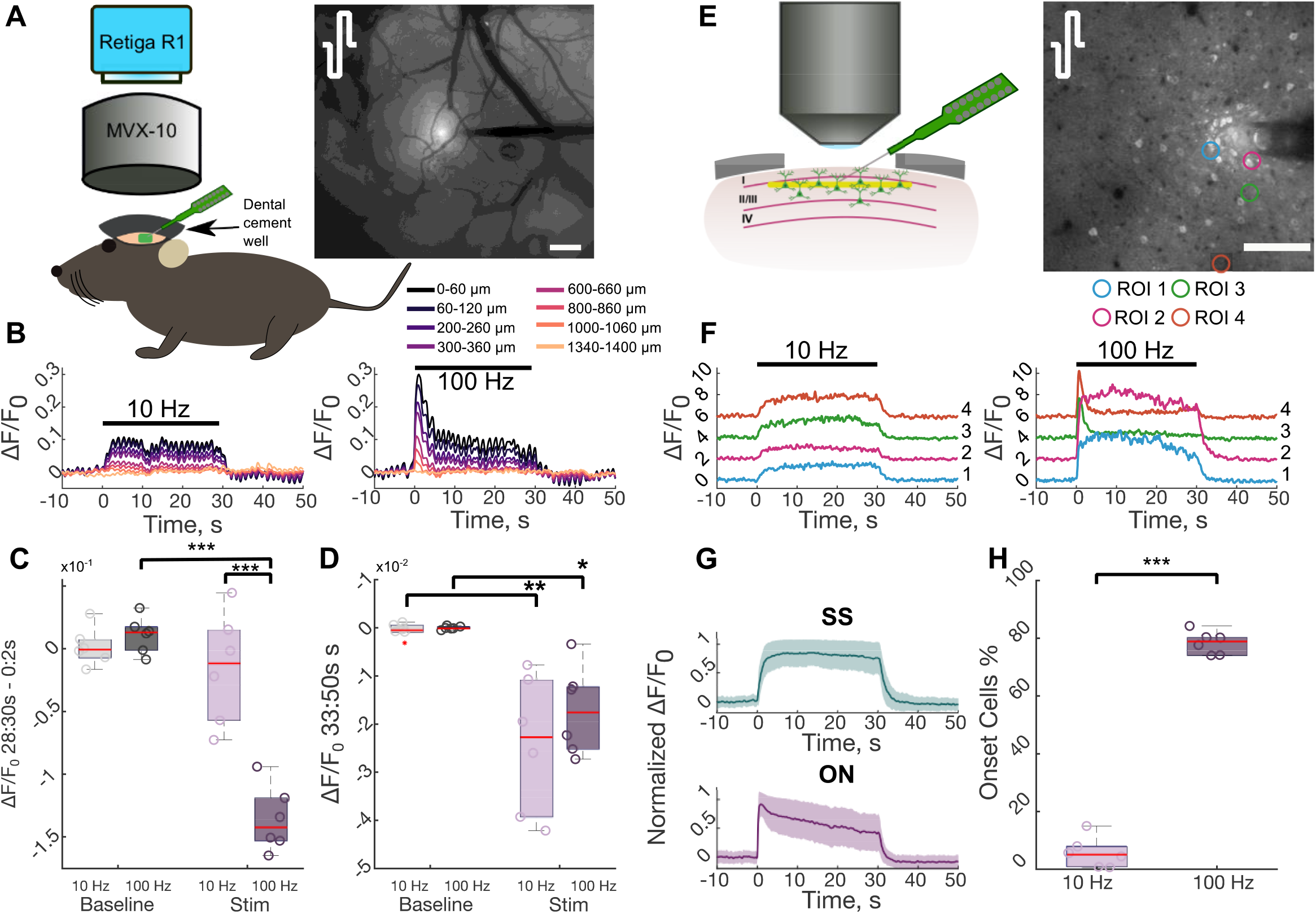
Calcium activity reduces over time during high frequency stimulation. **A**. An anesthetized Thy1-GCaMP6s mouse was implanted with an acute NeuroNexus electrode under mesoscale imaging to observe broader cortical activation during stimulation. The mean activation in during stimulation with symmetric cathodal-first pulses highlights that activity decreases with distance from the electrode. Scale = 200 μm. **B**. Calcium activity in radial bins around the electrode depicted in (A) reduces during 100 Hz-stimulation, in contrast to 10 Hz. **C**. 100 Hz-stimulation results in a significant reduction in stimulation-induced neural activity between the onset (0-2s) and the end (28-30s) of stimulation compared to 10 Hz-stimulation and the change in activity during between the beginning and end of the baseline period (2-Way repeated measures ANOVA Effects: Frequency, Stim/Baseline, Interaction: p<1e-3; post-hoc Welch’s T-Test). **D**. Post-stimulation neural activity is significantly depressed (below baseline) near the electrode compared to normal activity (2-Way repeated measures ANOVA Effects: stim/baseline: p<1e-4; post-hoc Welch’s T-Test). **E**. To observe stimulation induced neural activity at the cellular level, an electrode was implanted into layer II/III under a two-photon microscope. The standard deviation projection during 100 Hz-stimulation with symmetric cathodal-first pulses highlights the activated neurons. Scale = 100 μm. **F**. Stimulation induced neural activity in the regions of interest in (E) highlight that some neurons reduce their activity during 100 Hz-stimulation (right), whereas those same neurons display stable activation during 10 Hz-stimulation (left). **G**. Neurons that display reduced neural activity are defined as onset (ON) and neurons with stable activation over time are steady-state (SS). **H**. The majority of activated neurons reduce their activity during high frequency stimulation (Student’s T-test). *: p<0.05, ***: p<0.001.

### Stimulation

Stimulation was delivered via TDT RZ6D system controlling an IZ2 stimulator (Tucker-Davis Technologies, Alachua, FL) at 10 or 100 Hz using charge-balanced biphasic cathodal-or anodal-leading waveforms (Table 1). A monopolar setup was used to better isolate the waveform specific effects at a single electrode location(39). The asymmetry of the waveforms was defined by the ratio of the leading phase to the return phase(40) (Eqn. 1) and is comparable to waveforms that can be used clinically(4, 23). Stimulation trials involved a 30s baseline, 30s stimulation, and 20s post-stimulation period. All 8 stimulation waveforms were delivered in random order at 10 or 100 Hz for each session with a minimum of 5 minutes between trials.

**Table 1:** Parameters of stimulation waveforms used in this study. Waveforms are charge-balanced biphasic and were delivered at 10 or 100 Hz for 30s. All waveforms deliver 2.5 nC/phase, and a charge density of 0.36 mC/cm^2^.

| Asymmetry Index | Leading Amplitude, $\mu\text{A}$ | Leading Pulse Width, $\mu\text{s}$ | Return Amplitude, $\mu\text{A}$ | Return Pulse Width, $\mu\text{s}$ | Total Pulse Time, $\mu\text{s}$ |
| --- | --- | --- | --- | --- | --- |
| -1/2 | $\pm 12.5$ | 200 | $\mp 6.2$ | 400 | 600 |
| 0 | $\pm 12.5$ | 200 | $\mp 12.5$ | 200 | 400 |
| 4 | $\pm 2.5$ | 1000 | $\mp 12.5$ | 200 | 1200 |
| 7 | $\pm 1.6$ | 1600 | $\mp 12.5$ | 200 | 1800 |

All waveforms deliver 2.5nC/phase (0.36mC/cm^2^) which is within safety recommendations of microelectrodes(41, 42). Waveforms in figures are represented by the leading polarity with a subscript of the asymmetry index (e.g. C_0_ represents symmetric cathodal-leading).

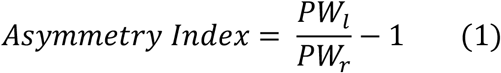

### Image analysis and statistics

#### Stimulation inclusion

Six biological replicates were obtained consistent with previous reports with sufficient statistical power(24, 28, 30, 36). Stimulation sessions, defined as one train of the 8 waveforms delivered at 10 or 100 Hz, were considered biologically independent when delivered in different locations (electrode sites or insertion location) as shown by Histed and colleagues(15). Sessions with significant vasculature displacement were excluded from analysis.

#### Neuronal soma and neuropil quantification

Neuron regions of interest (ROI) were manually outlined using a 2D projection of the standard deviation over the stimulation trial using ImageJ as previously done(24, 28, 29). The cellular location and mean fluorescence intensity of the ROI were then extracted for analysis in MATLAB and fluorescence was transformed into ΔF/F0 using the 30s pre-stimulus baseline. ΔF/F0 traces were then filtered (FIR cutoff frequency of 2Hz (24, 28, 36)) to remove artifacts.

Neurons were considered activated if ΔF/F0 exceeded the mean plus 3 standard deviations of the baseline activity for at least 1s (24, 28-30). Activity patterns were manually characterized based on the temporal dynamics of the calcium activity as either steady-state (SS) or onset (ON) activity similar to our previous reports(28, 30). Specifically, ON activity is defined as a rapid rise in calcium activity followed by a subsequent decrease through the stimulation train, whereas SS activity rises and remains elevated throughout the stimulation. The ‘activation time’ was calculated as the time it takes for the ΔF/F0 to reach half the maximum cumulative activation over the 30s stimulation train providing a metric to describe when the activity was focused (beginning or end)(30). For example, a SS neuron would have an activation time of ∼15s because the activity would be relatively evenly distributed across the stimulation train, while an onset neuron would have an activation time <15s because the activation is focused at the beginning of stimulation(30). Some neurons display greater calcium activity for one waveform over another, which we refer to as preferential activation. A neuron is defined to be preferentially activated by one waveform over another if the difference in magnitude of calcium activity during the last two seconds of stimulation between the two waveforms (28-30s) was greater than the average standard deviation during that period for the two waveforms (Eqn. 2).

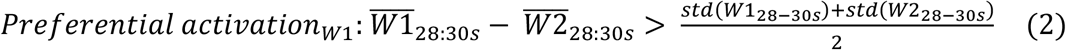

The neuropil, considered as mainly neuronal axons but also out of focus neuronal cell bodies and dendrites, can be independent of somatic activity(43-46). The neuropil and mesoscale calcium activity were determined in 20 μm radial bins around the electrode after masking out the electrode from the two-photon and mesoscale images(24, 28, 29). The identified neuronal ROIs were also masked out from the two-photon image. The average fluorescence in each bin was processed the same as the neuronal calcium activity. To determine the similarity of the neuropil activity with activated somata, the average ΔF/F_0_ in each two-photon neuropil bin was correlated with the calcium activity of neurons within the equidistant bin using Pearson’s Correlation Coefficient, R(28, 29).

#### Statistics

All statistical analyses were performed in R(47) with a significance level of 0.05. Analyses were grouped across biological replicates and consisted of N-way repeated measures ANOVA (ez package(48)) or a linear mixed-effects model (lme4 package(49)) followed by an F test. Regression analyses were performed using the lm function in R. When appropriate in the ANOVA, sphericity corrections were performed with the Greenhouse-Geisser method followed by Post-hoc pairwise unequal variance (Welch’s) T-tests with a Holm-Bonferroni correction of the alpha value. Error bars are standard error unless otherwise stated.

## Results

Mesoscale epifluorescence calcium imaging provides a network-wide view of neural activity, and two-photon imaging provides cellular resolution of neural activity. Using these two imaging modalities in a mouse model expressing the calcium sensor GCaMP6s in pyramidal neurons, (Fig. 1) we identified several patterns of ICMS-evoked activity: 1) sustained activity defined as suprathreshold calcium activity throughout a 30s stimulation train, 2) steady-state (SS) activity defined as a rise in calcium activity at the stimulation onset that did not reduce by the end of stimulation (*i*.*e*. stable), 3) onset (ON) activity defined as a rise in activity at the stimulation onset that substantially reduced by the end of stimulation but not necessarily below baseline, and 4) depression of activity defined as below baseline calcium activity that typically occurred after the end of stimulation. Because stimulation frequency and waveform asymmetry can influence the spatiotemporal activation of neurons around the electrode, we then tested whether these patterns changed between 10 and 100 Hz-stimulation and different waveform asymmetries.

### 100 Hz but not 10 Hz-stimulation results in substantial reduction of neural activity during stimulation

Cortical networks are thought to operate in an inhibition-stabilized network regime where feedback inhibition stabilizes excitatory network activity. This implies that feedback inhibition would reduce stimulation-induced neural activity after a short period of excitation(50, 51). Therefore, using symmetric cathodal-leading stimulation waveforms delivered at 10 or 100 Hz for 30s, we first tested whether stimulation-evoked activity reduced between the onset (0-2s) and the end (28-30s) of stimulation under mesoscale calcium imaging (Fig. 1A-D). Stimulation-induced activity was strongly elevated within 400 μm of the electrode and substantially reduced during 100 Hz stimulation (Fig S1A-B). Importantly, while 10 Hz-stimulation resulted in stable activity, 100 Hz-stimulation resulted in a 10% reduction in stimulation-induced activity within 400 μm of the electrode (Fig. 1B-C; 2-way ANOVA Effects: Frequency, Stim, Interaction p<1e-5). Interestingly, after the end of stimulation, both 10 Hz and 100 Hz-stimulation were followed by a long lasting (∼17s) period of depressed activity within 680 μm of the electrode (Fig. S1 C-D; Fig. 1C-D, 2-way ANOVA Effects: Stim p<1e-4).

Two-photon imaging of individual somata revealed a similar trend to mesoscale imaging, with “onset” activity in some neurons at 100 Hz, while others maintained “steady-state” calcium activity at both 10 Hz and 100 Hz (Fig. 1F-G). Notably, while only 5.8±5.3% of activated neurons displayed an ON response during 10 Hz-stimulation, the activity of 78.5±3.9% reduced during 100 Hz-stimulation (Fig. 1I, Welch’s t-test: p<1e-7) consistent with previous work(24, 28, 30). Although reduced entrainment probability over time could explain the reduction in stimulation-induced activity(52, 53), the stable activity of SS neurons suggest that some cells do maintain the same level of entrainment throughout the train. Therefore, SS responses (Fig. 1H) are thought to be due to direct, antidromic activation, while the ON responses (Fig. 1H) are thought to be due to indirect, trans-synaptic activation followed by inhibition(28, 36) (See Discussion (28)). Altogether these data suggest inhibition may contribute to the progression of network activity during 100 Hz-stimulation, but may not exceed stimulation-induced excitation during 10 Hz stimulation.

Electrical activation of neurons depends on the geometrical and electrical properties of neural elements (soma, axon dendrite), which differs between subtypes of excitatory and inhibitory neurons(10, 11, 14, 54, 55). Modeling studies predict that cathodal-first waveforms with a longer leading phase relative to the return phase (*i*.*e*. more asymmetric) have greater selectivity of activation at the soma, while asymmetric anodal-first waveforms are more selective for activation of passing fibers compared to symmetric waveforms(14). Therefore, we hypothesized that the reduction in stimulation-induced mesoscale neural activity at 100 Hz would depend on waveform asymmetry (Fig. 2, Table 1). Consistent with our previous study demonstrating that waveform asymmetry modulated spatial activity of overlapping populations of cortical neurons at 10 Hz-stimulation (24), waveform asymmetry also modulated the spatial activation of cortical neurons at 100 Hz (Fig. 2B). However, stimulation-induced neural activity within 400 μm reduced by ∼6-12% during 100 Hz-stimulation. This reduction was significantly greater than the difference in activity between the first and last two seconds of the baseline period (Fig. 2C-D 3-way ANOVA Effects: Stim, Asymmetry-Polarity interaction: p<0.01). Additionally, the reduction in calcium activity during ICMS depended on waveform asymmetry with highly asymmetric (index of 7) cathodal-first pulses resulting in significantly less reduction in induced activity over the pulse train than anodal-first highly asymmetric pulses. Notably, all waveforms resulted in comparable post-stimulation depression of calcium activity (Fig. 2E-F 3-way ANOVA: Stim: p<1e-4) independent of frequency (Fig. 3B), which suggests that the mechanisms mediating the reduction during stimulation may be different than those after the pulse train. Importantly, the reduction in activity during and after the 100Hz pulse train correlated with the magnitude of activation at the onset (Fig. 2G-H; Fig. 3 C-D) and was greater than the reduction at 10 Hz for all waveforms (Fig. 3A 3-way ANOVA; Frequency, Asymmetry, Stim: p<0.01). Together, these results suggest that waveform asymmetry and frequency modulate the activity at the onset of stimulation and the reduction of stimulation induced activity. Notably, however, the waveform asymmetry and polarity can modulate the magnitude of stimulation-induced calcium activity(24). Therefore, because the reduction in stimulation-induced activity correlated with the of activation at the onset (Fig. 2G, Fig. 3C), the waveform-dependent reduction in stimulation-induced activity (Fig. 2D) may be a result of the previously described waveform asymmetry differences in the magnitude of activation rather than waveform-dependent activation of different populations.

**Figure 2.**
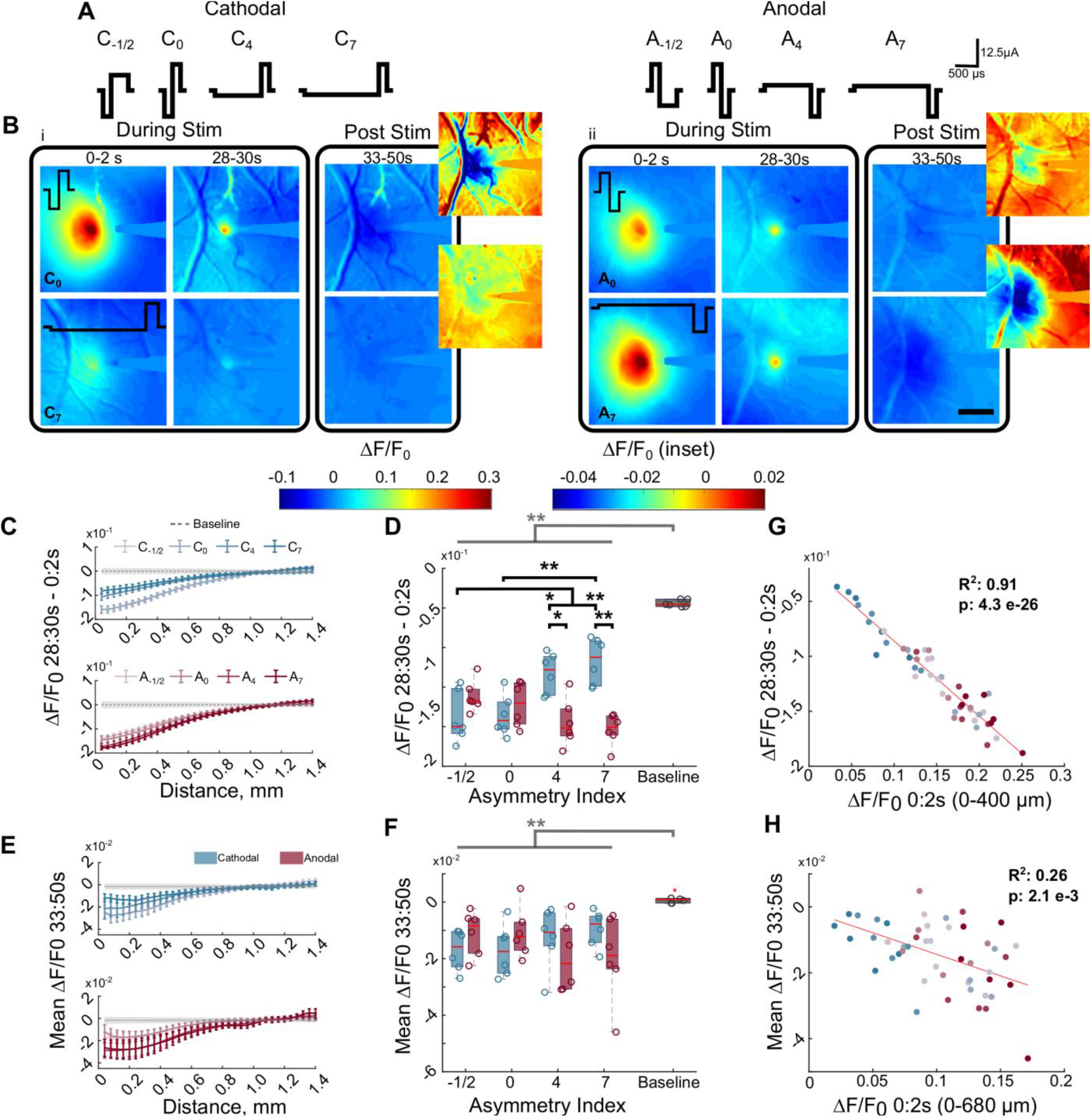
Neural activity reduces over time for all waveforms of stimulation when delivered at 100 Hz. **A**. Graphical representation of all waveforms tested. The asymmetry index increases as the duration of the activation phase increases relative to the return phase. Control waveform: cathodal-first symmetric with 200 μs phase duration, 12.5μA and 2.5 nC/phase. **B**. The magnitude of neural activation and the reduction in neural activity over time depends on the stimulation waveform and polarity. Symmetric cathodal-first (i, top; C_0_) and asymmetric anodal-first waveforms (ii, top; A_7_) have greater magnitude of activation and reduction during stimulation in neural activity compared to asymmetric cathodal-first (i, bottom; C_7_), and symmetric anodal-first (ii, bottom; A_0_), respectively. Following stimulation (Post Stim) there is a long-lasting (∼17s) depression of activity below baseline (dark purple/black) Scale bar: 500 μm. **C**. The reduction in neural activity from the onset and end of stimulation reduces with distance from the electrode. Grey represents the mean baseline activity ± the average standard deviation during baseline. **D**. All waveforms of stimulation display a greater reduction in neural activity over time during stimulation compared to normal changes in activity during baseline (3-way rmANOVA, Effects: stim/baseline, Polarity-asymmetry interaction p<0.01). The change in baseline activity was calculated as the difference in calcium activity between the first and last two seconds of the baseline period. **E**. 100 Hz-stimulation is followed by a long-lasting (averaged over 3-20s post-stimulation) and wide-spread depression of neural activity focused near the electrode. Grey represents the mean baseline activity ± the average standard deviation during baseline. **F**. The post-stimulation activity is significantly reduced relative to normal activity (baseline) over 680 μm for 100 Hz (2 way and 3 way -repeated measures ANOVA). Baseline activity was calculated as the mean activity during the last 17s of the baseline period. **G**. Greater activity at the onset of stimulation (0-2s) is significantly correlated with greater reduction by the end of stimulation (28-30s). **H**. Greater activity at the onset of stimulation (0-2s) is significantly correlated with greater post-stimulation depression of activity (3-20s post stimulation) within 680 μm of the electrode. *: p<0.05, ** p<0.01. Grey bar and * indicate main effect from Stim vs Baseline

**Figure 3.**
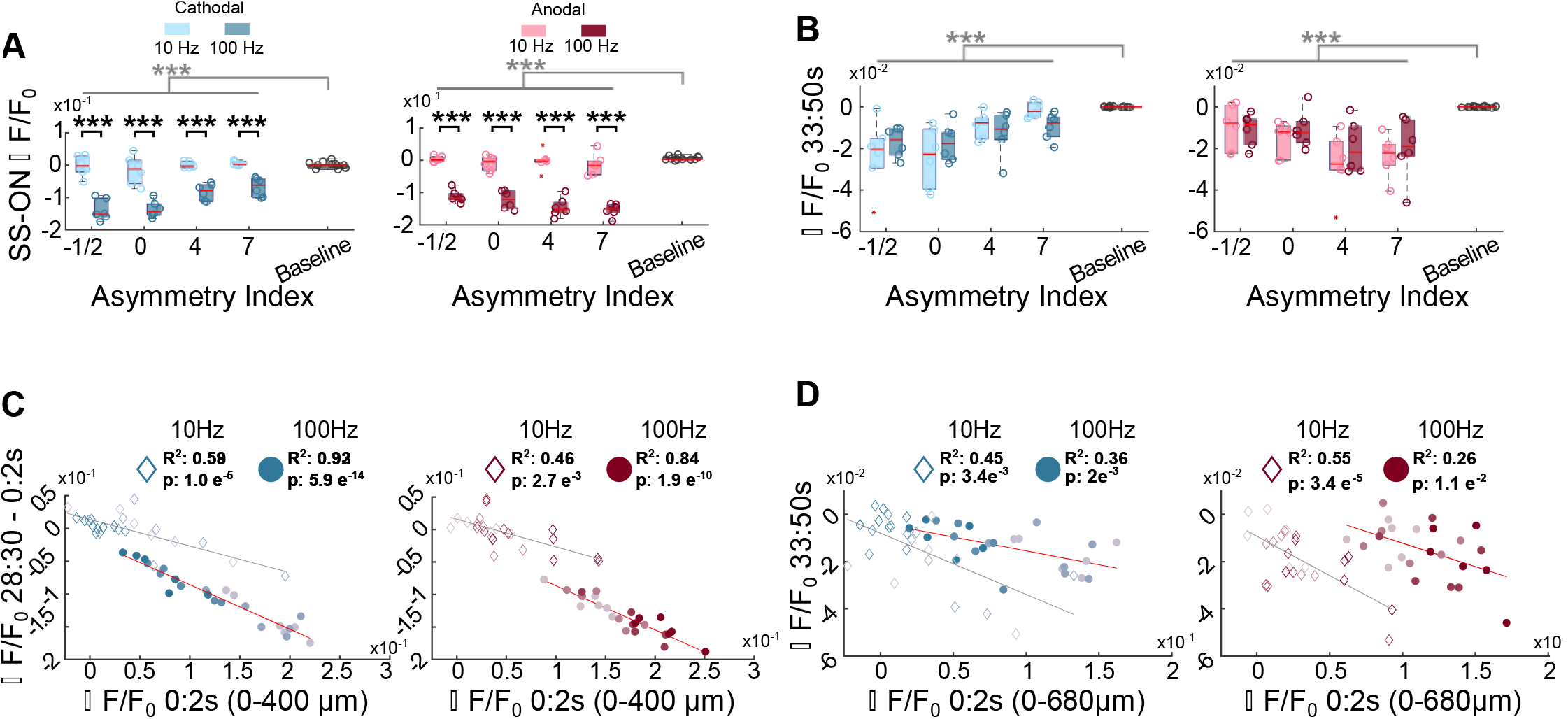
Trends in neural activity are consistent across waveforms between different frequencies. **A**. 100 Hz-stimulation results in a greater reduction in neural activity than 10 Hz-stimulation for all waveforms and baseline (3-way repeated measures ANOVA). **B**. Stimulation is followed by a period of depression below baseline following stimulation for all waveforms and both 10 Hz and 100 Hz-stimulation (3-way ANOVA). Gray bars represent significant main effect between stim/baseline in 3-way ANOVA. **C**. The magnitude of stimulation induced activity at the onset correlates with the reduction in stimulation induced activity for 10 Hz (diamonds) and 100 Hz (circles). **D**. The magnitude of stimulation induced activity at the onset correlates with the magnitude of the post-stimulation depression of activity for 10 Hz (diamonds) and 100 Hz (circles) ***: p<0.001.

### Two-photon imaging during 100 Hz-stimulation demonstrates waveform alters the pattern and magnitude of direct (SS) or indirect (ON) activation around the electrode

Modeling studies suggest that stimulation waveform may activate different populations of neurons depending on polarity and asymmetry(13, 14); therefore, we next used two-photon imaging to evaluate how waveform asymmetry influenced the population-level neuronal recruitment. We first hypothesized that stimulation waveform would modulate the population of neurons that were directly or indirectly activated (Fig. 4). We tested whether stimulation waveform asymmetry could predict if a neuron had an SS response (direct) or an ON response (indirect) during 100 Hz stimulation. 100 Hz stimulation was used because ON responses are rarely observed during 10 Hz stimulation (Fig. 1H)(30). While 72-86% of neurons displayed an ON response (Fig. 4 A-E), not all neurons that display an SS response for one waveform have the same response for all waveforms. Notably, only 18% of all activated neurons solely responded as SS independent of waveform (Fig. 4F), and these neurons were concentrated within 40 μm of the electrode (Fig. 4G). This is consistent with previous data suggesting that direct activation of neurons or neuronal elements occurs within a small region around the electrode as shown by stable glutamate release in the same area(28).

**Figure 4:**
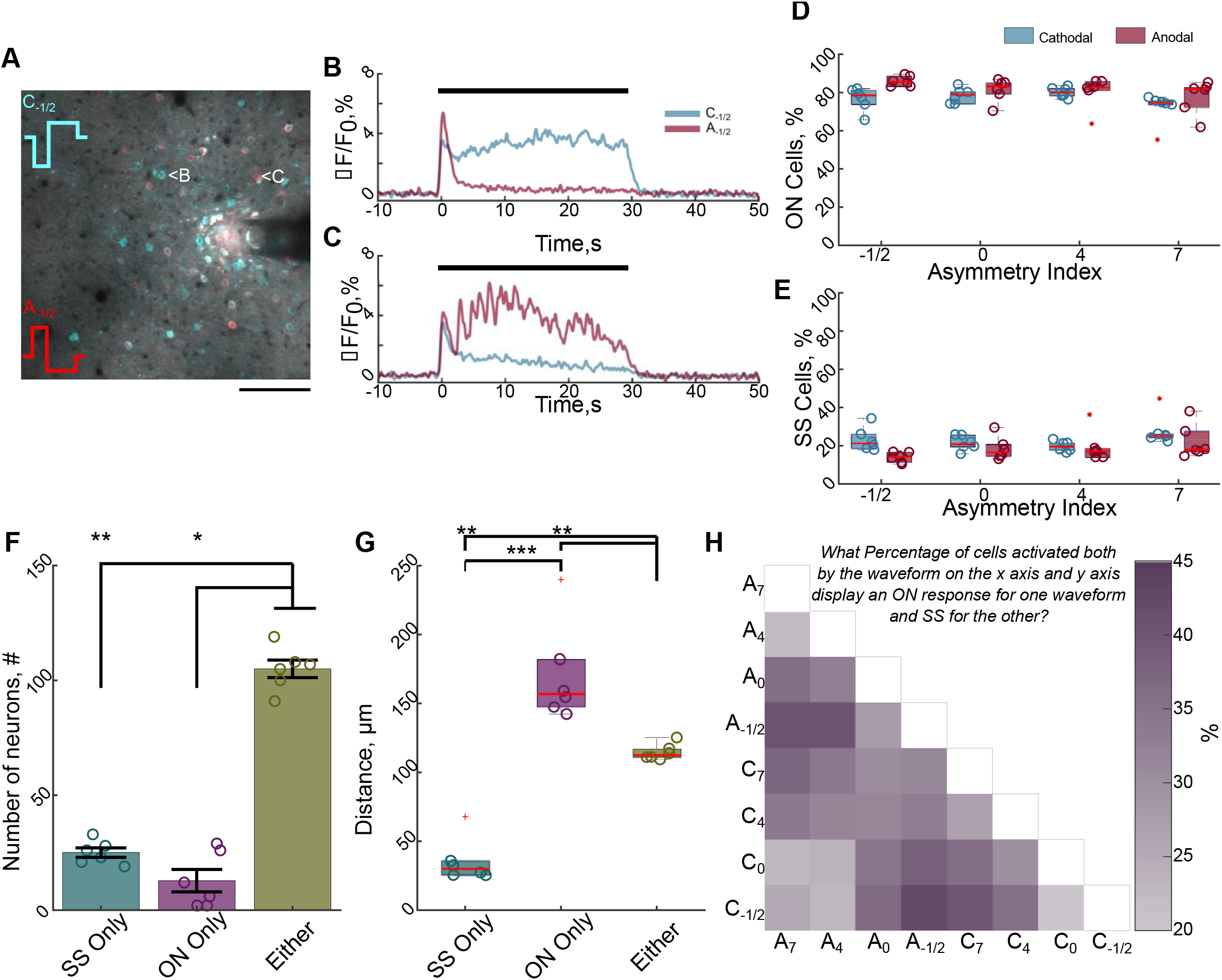
Stimulation waveform modulates which population of neurons are directly and indirectly activated. **A**. Different populations of neurons display greater or more sustained activation for different waveforms demonstrated by a composite image of the 2D projection of the standard deviation during stimulation with a cathodal (cyan) or an anodal (red) waveform. Neurons with greater cyan color have greater or more sustained activation for the cathodal waveform, whereas red neurons have greater or more sustained activation for the anodal waveform. White neurons have similar magnitudes of activation for both waveforms. Scale bar: 100 μm. **B-C**. The calcium traces in of the neurons depicted in (A) during cathodal or anodal-first waveforms reveal that waveform can modulate whether a neuron is directly activated (stable) or indirectly activated (reduced activity over time). Black bar indicates stimulation time. **D-E**. The majority of activated neurons display an onset response (reduced activity over time) independent of waveform. **F**. Only ∼18% of all activated neurons responded as steady-state (SS) independent of waveform, suggesting only a small percentage of neurons are directly activated (Kruskal-Wallis, p<0.01; post-hoc Dunn’s Test). **G**. The directly activated neurons (SS) are located within 40 μm of the electrode on average (ANOVA: p<1e-5; post-hoc Welch’s T-test). **H**. Waveforms expected to activate different populations of neurons (*e*.*g*. symmetric cathodal (C0) vs asymmetric cathodal (C7)) are more likely to result in a greater percentage of neurons being directly activated (SS) for one waveform and indirectly activated (ON) for another. Darker purple indicates a greater percentage of neurons activated by both waveforms responded as SS for one waveform and ON for the other. *: p<0.05; **: p<0.01; ***:p<0.001.

Moreover, because modeling studies suggest waveforms with different asymmetries are more selective for different neural populations(13, 14), we would expect similar waveforms would predict a similar response type (*e*.*g*. SS response for two similar waveform asymmetries). We therefore hypothesized that increasing the difference in asymmetry index would raise the probability that a given cell would change its response profile (*e*.*g*., SS for one waveform, ON for another). In fact, larger differences in the waveform asymmetry were twice as likely to engender a change in the response type (Fig. 4H dark purple e.g. C_-1/2_ – C_7_ = 39.8%) compared to waveforms with similar asymmetries (Fig. 4H light purple e.g. C_-1/2_ – C_0_= 20.4%). This suggests that waveform may be changing which populations of neurons are directly or indirectly activated. Although, it is also possible that waveform is modulating activation and/or entrainment of different excitatory and inhibitory neurons(13, 52, 53).

Because stimulation waveform influences the response profile of different neurons (Fig. 4), it is possible that subpopulations of these excitatory neurons are more strongly recruited by certain waveform asymmetries. Therefore, we determined which neurons were preferentially activated (Eqn. 2) for one waveform over another (Fig. 5). While some neurons that were activated in common between two waveforms had similar magnitudes of activity (Fig. 5A-D; white), 20-90% of the neurons activated were preferentially activated for at least one waveform over another (Fig. 5E). Importantly, although some waveforms generally elicit greater activity (e.g. symmetric cathodal C_0_, asymmetric anodal A_7_)(24, 56) a small percentage of neurons prefer waveforms that are expected to require higher amplitudes to elicit the same amount of activity(14, 40) (e.g. C_4_, C_7_; Fig. 5E, Fig. S2). This suggests certain neural populations may be preferentially recruited with specific waveforms(14).

**Figure 5:**
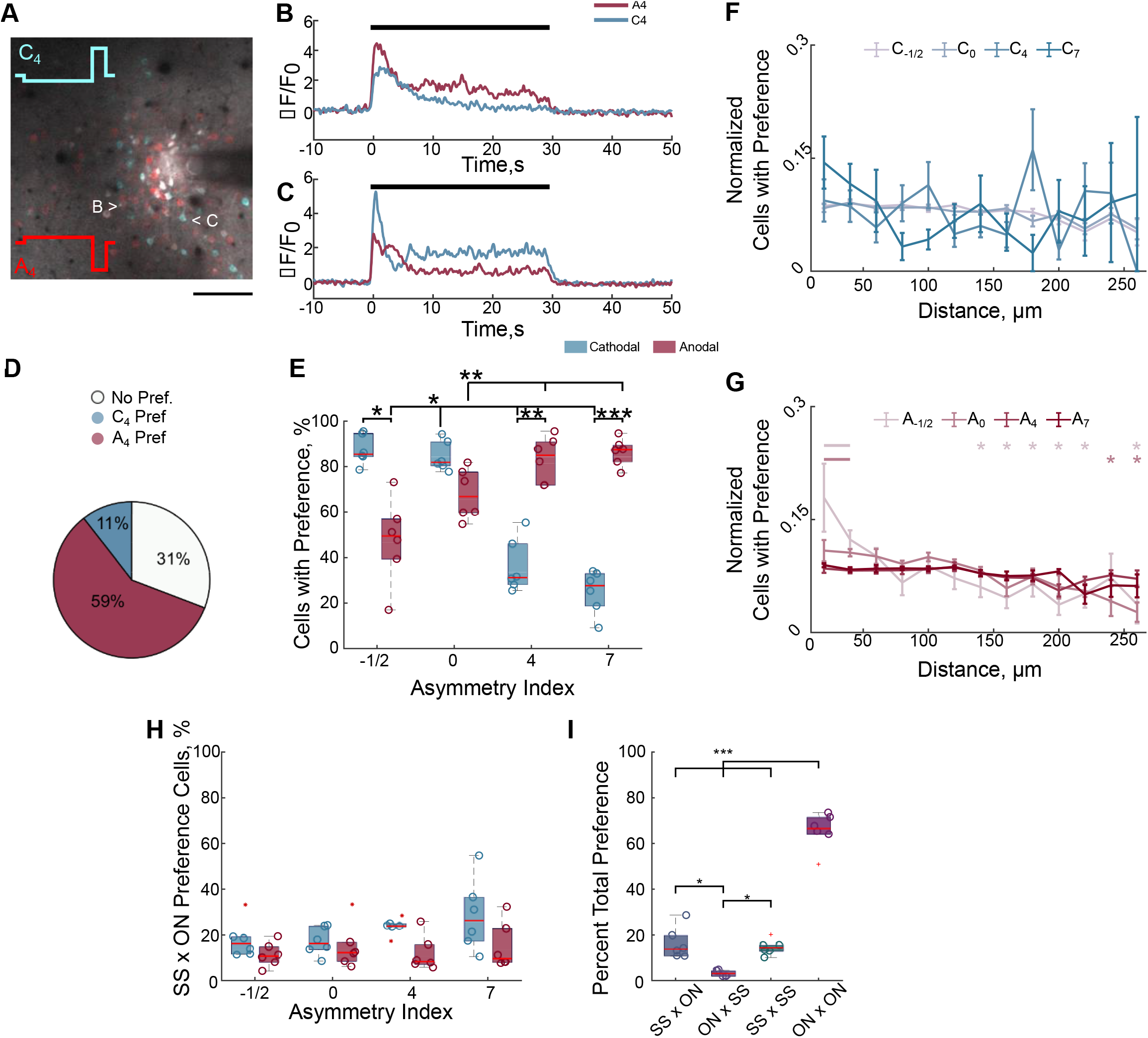
Stimulation waveforms elicit greater activation in certain neural populations over others during 100 Hz. **A-D**. Certain neural populations are preferentially activated by cathodal-first asymmetric waveforms with long leading phases (Cyan neurons, 11% of neurons activated in common), while others are preferentially activated by anodal-first waveforms with the same asymmetry (Red; 59% of neurons activated in common). Neurons with similar magnitudes of activation between the two waveforms are white (31%). Preferential activation is defined in Eqn 2 of the methods. Scale 100 μm. **E**. Certain populations of neurons are preferentially activated by different waveforms over at least one other waveform. The proportion significantly depends on polarity and asymmetry (2-way repeated measures ANOVA, Polarity, Asymmetry, interaction p<0.01; post-hoc Welch’s T-Test). **F-G**. Preferential activation is generally distributed over distance for cathodal-first waveforms (ANOVA run on linear mixed model Distance p>0.05), while some neurons within 40 μm of the electrode are more likely to be preferentially activated by some anodal-first waveforms than more distal neurons (ANOVA run on linear mixed-model: Distance, Asymmetry-Distance interaction p< 0.001; post-hoc Welch’s T-test). Data were normalized to the number of cells at each distance and to the total over distance. **H**. There is a low probability of preferential activation being due to the fact that some neurons are directly activated (SS response) for the preferred waveform and are indirectly activated for the other (ON response; 2-way rm ANOVA Polarity p<0.05). **I**. The majority of preferential activation occurs when a cell is indirectly activated by both waveforms (ON x ON), whereas preference is similarly likely to occur when the neuron is directly activated by both waveforms (SS x SS) or if the neuron is directly activated by the preferred waveform and indirectly for the other (SS x ON). It is not likely that a neuron is preferentially activated by a waveform that elicits activity through indirect activation when another waveform activates the neuron directly (ON x SS). Data were pooled across all waveform comparisons *: p<0.05, **:p<0.01, ***:p<0.001

Because some waveforms may preferentially elicit activity in proximal neurons (C_4_, C_7_), while others would activate distal neurons (A_4_, A_7_)(14, 24), we hypothesized that waveform preference would depend on distance from the electrode. Therefore, we quantified the percent of activated neurons displaying preferential activation in different radial bins and normalized to the total percent of cells with preferential activation. Surprisingly, distance generally did not influence preferential activation (Fig. 5F-G, linear mixed model) suggesting a more complex recruitment pattern.

Additionally, preferential activation could depend on whether the neuron was directly activated by one waveform but not the other (SS vs ON; Fig. 4B-C). However, only 18-28% of preference was due to a neuron displaying an SS response for the preferred waveform and ON for the other (SS x ON preference; Fig 5H). In fact, the majority of preference occurred when the neuron was indirectly activated during both waveforms (Fig. 5I ON x ON; 66%). There was similar frequency of preferential activation when the neuron was directly activated by both waveforms (Fig. 5I, SS x SS, 15%) or when the neuron was directly activated by the preferred waveform and indirectly by the other (Fig. 5I, SS x ON, 16%). In rare occurrences (3%) indirect activation resulted in greater calcium activity than direct activation (Fig. 5I, ON x SS). This suggests that the preference for one waveform over another is largely due to differences in the cumulative network activation driving greater indirect activation in some neurons (ON x ON). However, preference is also substantially influenced by stronger direct activation (SS x SS), or direct activation outweighing indirect activation (SS x ON). Altogether, these results suggest that waveform can modulate which neural populations are preferentially recruited through direct and indirect activation.

### Waveform and frequency modulate the spatial and temporal activation of neurons around the electrode

Is unclear how waveform asymmetry-dependent activation of overlapping neural subpopulations influences the progression of activity over time at different frequencies. Therefore, we next tested if waveform and frequency differentially modulate spatial activation of neurons and the progression of population activity over time (Fig. 6). The greatest differences in the magnitude of activation between 10 Hz and 100 Hz-stimulation is concentrated within 40 μm of the electrode (Fig. 6A-B, linear mixed model Polarity, Frequency, Distance: p<0.01). Additionally, the distribution of SS cells is significantly elevated within the first 60 μm for both frequencies (Fig. 6C; 3-way ANOVA: Frequency, Distance p<0.05), though the density is elevated for 100 Hz. Interestingly, the density of ON cells initially increases from 20 to 60 μm before decreasing for 100 Hz-stimulation, but increases with distance for 10 Hz (Fig. 6D).

**Figure 6.**
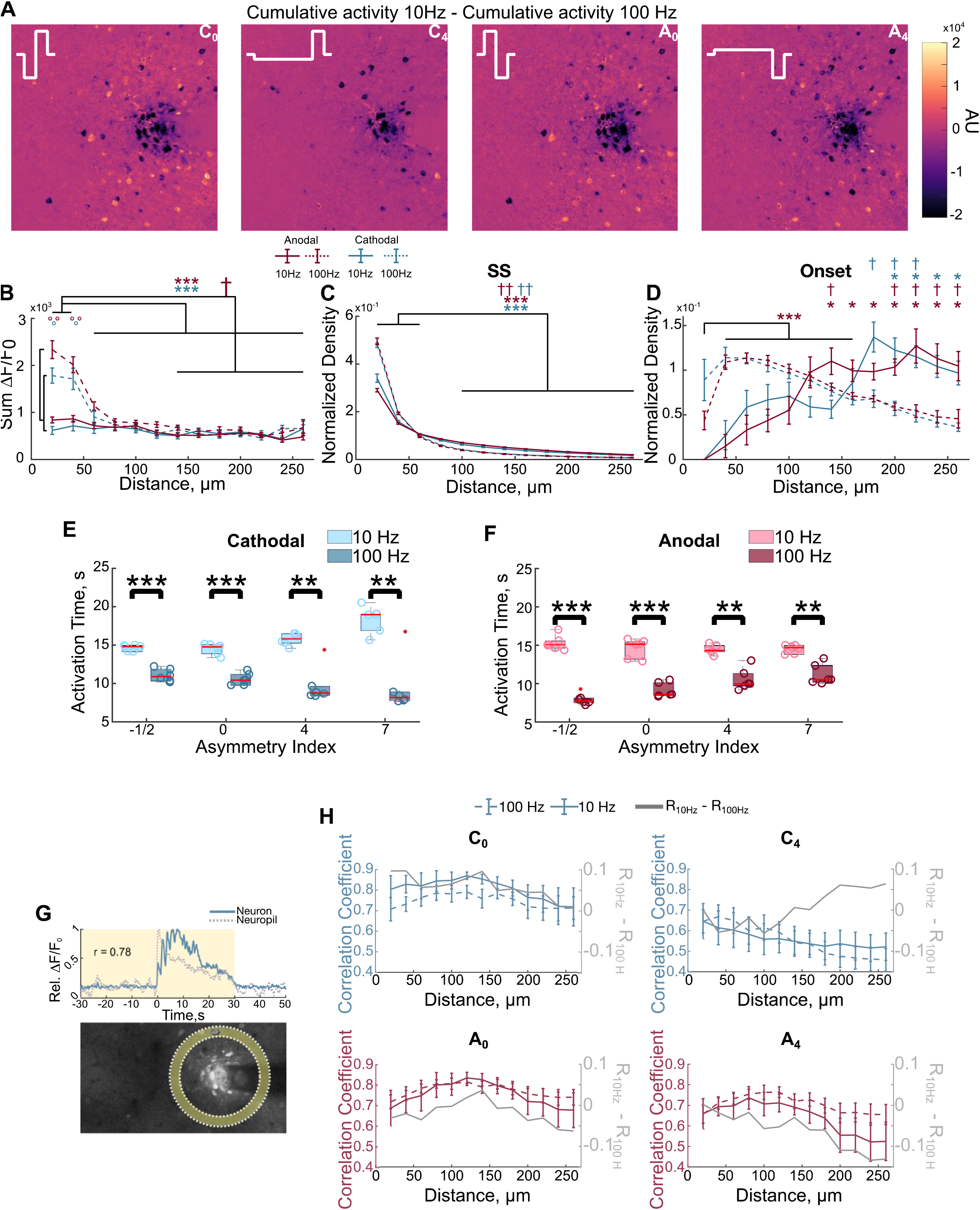
Stimulation waveform and frequency differentially modulate the neuronal and neuropil activity around the electrode. **A**. The difference between the cumulative activity at 10 Hz and 100 Hz highlights that the greatest difference in activity is concentrated close to the electrode. **B**. Neurons are most strongly activated within the first 40 μm and is significantly elevated for 100 Hz and quantified by the cumulative fluorescence during stimulation (F-Test on linear mixed model Polarity, Frequency, Distance: p<0.01; post-hoc Welch’s T-test). **C**. The density of steady-state cells, normalized to the density of activated neurons and total density is concentrated within the first 60 μm, but is more distributed for 10 Hz-stimulation (3-way ANOVA: Frequency, Distance p<0.05). **D**. The density of onset cells is more distributed, but follows a complex spatial pattern during 100 Hz. Significance is with respect to the first 40 μm (3-way ANOVA, Welch’s T-test post hoc, Bonferroni-Holm correction), with the exception of anodal 100 Hz due to the significant increase from 20μm to 40-160μm. **E-F**. The calcium activity of neurons is greater in the first half of stimulation during 100 Hz. This effect is greater for some waveforms as quantified by the activation time. **G**. Representation of the calculation of the neuron-neuropil correlation. Pearson correlation was computed between neuronal calcium activity with the calcium activity of the neuropil bin at the same distance. **H**. The correlation between neuronal and neuropil activity follows different spatial patterns between 10 Hz and 100 Hz depending on the stimulation waveform (2-way ANOVA: C_0_-Distance p<1e-7; C_4_-Distance p<1e-15, interaction p<0.01; A_0_-Distance p<1e-13; A_4_-Distance p<1e-9) *: p<0.05, **: p<0.01, ***: p<0.001. * in the respective color of anodal (red) or cathodal (blue) indicate significance for 100 Hz. † in the respective color indicates significance for 10 Hz. ° in the respective color indicates significance between 10 Hz and 100 Hz.

We next quantified the time it takes for a neuron to reach half of its cumulative calcium activity during the pulse train (*i*.*e*. activation time(30)) to test if differential direct and indirect activation of neurons modulates the progression of population activity over time. For example, activity focused at the beginning or end of stimulation will have an activation time <15s or >15s, respectively. Interestingly, as asymmetry increased for cathodal-first waveforms (or decreased for anodal-first) population activity tended to increase over time during 10 Hz-stimulation, but decreased to a greater extent during 100 Hz-stimulation (Fig. 6E-F).

Together these data suggest that direct activation of axons and dendrites occurs within 40-60 μm of the electrode, and trans-synaptic recruitment of neurons has a more complex spatial pattern. It is unclear how these patterns lead to the observed waveform asymmetry and frequency-dependent activity over time (Fig. 6E-F). In calcium imaging, the neuropil has been suggested to largely consist of directly activated axons(43). Therefore, to test if the progression of population activity depended mainly on the pattern of direct activation, we next quantified correlation of neuronal calcium activity with that of the neuropil bin at the same distance (Fig. 6G). If the activity in the axons of directly activated excitatory neurons were solely responsible for the temporal progression of the somatic activity of the neural population, we would expect to see a trend in the neuron-neuropil correlation that is similar for waveforms that produce similar temporal progression of activity (*e*.*g*. C_4 &_ A_0_ vs C_0 &_ A_4_). In contrast, the difference in the neuron-neuropil correlation between 10 Hz and 100 Hz followed different patterns for these waveforms. Specifically, the correlation was larger during 100 Hz for many waveforms (C_4_, A_0_, A_4_) and became less than that of 10 Hz with distance for one waveform (C_4_), or became even larger with distance for others (A_0_, A_4_). These results suggest, the resulting population activity depends on more than the cumulative activity of excitatory axons (*e*.*g*. subpopulations of inhibitory neurons), which supports a need for further investigation into the mechanisms of intracortical microstimulation and contributions of other cell types.

## Discussion

It remains unclear which neural populations around the electrode are activated and how stimulation parameters can modulate their activity to selectively elicit specific artificial percepts during ICMS. We previously demonstrated that 10 Hz ICMS waveform asymmetry modulates the magnitude and the spatial localization of neuronal activation(24). In this present study, we tested if stimulation waveform asymmetry modulates the pattern of recruitment (direct vs indirect) of different populations of cortical pyramidal neurons and how those differences shape stimulation-induced population activity. Delivering asymmetric stimulation waveforms at 100 Hz demonstrated that the waveform impacts the stability of activation (SS vs ON) of different, but overlapping populations of neurons (Fig. 4), suggesting that waveform modulates the pattern of directly activated neurons. Importantly, neurons within 20-60 μm of the electrode displayed strong and stable activation independent of stimulation waveform (Fig. 4, Fig. 6) which supported our hypothesis that direct activation of somata and axons/dendrites occurs within this region(28, 29). Furthermore, certain neuronal populations responded with greater calcium activity to specific waveforms (Fig. 5, Fig. S2), which was not simply a result of direct vs indirect activation. Moreover, the progression of population calcium activity was differentially modulated over time by frequency and waveform asymmetry (Fig. 6), suggesting that the resultant network activity is a combination of preferential activation of different neuronal populations by waveform and frequency. Ultimately, these results highlight the complexity of neuronal activation and may have important implications for using different waveforms to modulate perception in neural prosthetics.

### Stimulation waveform modulates neural recruitment patterns

Understanding how different stimulation parameters influence activation of neuronal populations could help inform stimulation paradigms for sensory restoration. Importantly, we demonstrate that certain populations of neurons are preferentially recruited with different waveform asymmetries (Fig. 5, Fig. S2). Consistent with other reports(24, 40, 57-59), cathodal-first symmetric pulses and highly asymmetric (index of 7) anodal-first pulses elicited larger activity than highly asymmetric cathodal-pulses (Fig. 2, 5, S2.). This effect correlated with the amount of reduction in activity during the ICMS train, and the magnitude of the post-stimulation depression in activity (Fig. 2). Because waveforms that elicit larger magnitudes of neural activation also increase the number of neurons activated, it is possible that the larger reduction in neural activity is due to recruitment of more inhibitory neurons(24). Despite this main effect, some sub-populations of neurons were preferentially activated with waveforms that generally elicit less activity (e.g. A-1/2,C7 Fig. 5, S2). Although these populations were small, the subtle differences in population activation by these waveforms consistently drove unique progression of population activity in a frequency-dependent manner (Fig. 6E-F), which could not be fully explained by patterns in the correlation between activated neurons and the neuropil (Fig. 6H). Therefore, due to the electrophysiological differences between neuronal subtypes (i.e. excitatory inhibitory) and neuronal elements (i.e. axon, soma, dendrites), it is possible that the waveform-dependent preferential activation of certain neuronal populations is a result of both the activation threshold as well as entrainment probablility for specific waveforms. For example, because inhibitory neurons are sparsely distributed and have distinct morphology, electrophysiological properties, and influence on excitatory neuron activity, it is possible that the proximity to different inhibitory neurons is responsible for these waveform and frequency differences (Fig. 6)(55, 60-62).

Additionally, there was asymmetry-dependent switching of the response type of neurons during 100 Hz stimulation that could not be observed during 10 Hz stimulation. Specifically, a neuron could respond as SS for one waveform and ON for another (Fig. 4). This suggests that although the magnitude of activation is greater for some waveforms, the stimulation waveform likely modulated which populations were being directly or indirectly activated, which could be important for downstream circuit responses. The mechanisms governing the “onset” effect have not been fully elucidated (Fig. 1-4); however, it is thought that these neurons are activated trans-synaptically and inhibition overcomes the synaptic excitation(28). It is also possible that the activity reduction during stimulation is due to virtual anode formation(11, 63, 64), entrainment failure(43, 65), frequency-dependent entrainment probability(13, 52, 53), or even neurotransmitter depletion(66). Importantly, the neuropil activation and glutamate release are stable during ICMS(28, 29) and SS responses are still observed farther (>60 μm) from the electrode(30), suggesting that inhibition is the main contributor to the reduction in activity.

It is also important to consider that not all SS neurons are directly activated. For example, because somatostatin (SST) neurons inhibit both excitatory and other inhibitory neurons, the stable calcium activity could be due to disinhibition(55). While pharmacological blockers of synaptic transmission could further dissect this mechanism, the dense, stable, and strong recruitment of SS neurons close to the electrode (Fig. 4, 6), their elevated correlation with the neuropil compared to ON neurons(28), and the observation of sustained glutamate release(28, 29) highly suggest direct activation of neurons and passing axons/dendrites is occurring close to the electrode, consistent with Histed and colleagues(15). Furthermore, the fact that SS responses only accounted for a small percentage of the activated neurons during 100 Hz stimulation (Fig. 1, Fig. 4) is consistent with a previous report suggesting that layer 5 neurons are mainly activated trans-synaptically during ICMS(67). Future studies could also take advantage of optogenetics to selectively silence inhibitory neuron populations(68) to help determine their role in reducing the activity of certain neuron populations during ICMS. Altogether, these data expand our current understanding of ICMS by highlighting how stimulation waveform influences the pattern of activation of different populations of excitatory neurons and how subtle differences in population recruitment may influence the progression of population activity.

### Implications for neural technology

There is substantial variability in the elicited percepts during ICMS(1, 6, 12). For example, a human participant reported that some electrodes evoked “tapping” sensations at 20 Hz and “buzzing” sensations at 100 Hz, while other electrodes evoked “tingle” at 100 Hz and no sensation at all at 20 Hz.(6). Because the number of neurons activated(69), their spiking pattern (70) and role in excitatory or inhibitory subnetworks(18, 71-74), may all modulate perception, even small changes in the population of neurons recruited (Fig. 4-5, Fig. S2) may have substantial effects on artificial percepts. Moreover, the specialized geometry and excitability of inhibitory neurons may predispose them to preferential activation by different waveforms, frequencies, or temporal patterns(13, 53, 61, 71). Therefore, waveform asymmetry could represent another dial to tune population activity and perception especially when there is variability between individuals(75) and electrodes within an individual(6). More specifically, a better understanding of how different stimulation parameters (e.g. waveform asymmetry, frequency, temporal pattern) modulate stimulation-induced population activity could aid the design of paradigms to selectively elicit specific percepts, such as pressure, across different electrodes despite differences in neural composition around the electrode sites.

Adaptation to stimulation, or a desensitization to a sustained stimulation, is a common feature of sensory systems may be beneficial for neural prosthetics(76). The magnitude of perceived artificial percept has been shown to decrease during continuous electrical stimulation of the somatosensory cortex(77), visual cortex(78), and peripheral nerves(76). This effect has been shown to be frequency-dependent, where increasing the frequency from 100 Hz to 300 Hz increased the rate of adaptation and adaptation did not occur during 20 Hz stimulation. Therefore, it is possible that the reduction in neural activity observed during 100 Hz stimulation (Fig. 1, 2) could be perceived as a reduction in the magnitude of the artificial sensation. Additionally, it is possible that the waveform asymmetry-dependent population responses that result in shorter or longer activation times (Fig. 6E-F) could result in faster adaptation or increases in the perceived magnitude, respectively, which could be beneficial for encoding perception magnitude changes during different tasks involving bionic prostheses. However, It should be noted that long-lasting depression of neural activity followed stimulation (Fig. 1-3), reminiscent of stimulation induced depression of neural excitability (SIDNE)(79-81). This may lead to reduced perceptual sensitivity to ICMS when delivered with short latency between stimulation(77). This effect is unlikely to significantly impact the results presented here because stimulation waveforms were delivered in random order with a minimum of 5-minute recovery time. Altogether, these data highlight the many factors contributing to the spatial and temporal activation of cortical excitatory neurons during ICMS (polarity, asymmetry, frequency) and suggest that future studies should investigate the activity of both excitatory and subtypes of inhibitory neurons to understand how stimulation waveform could increase the selectivity and specificity of eliciting artificial percepts.

In addition to sensory restoration with neural prosthetics, electrical stimulation is used clinically to treat symptoms of neurological disorders, such as Parkinson’s Disease(82). A recent study demonstrated that deep brain stimulation (DBS) could achieve long-lasting therapeutic benefit in a mouse model of Parkinson’s Disease through population-specific electrical neuromodulation(26). Importantly, by observing progression of stimulation-induced activity over longer stimulation trains (30s), this study highlighted that electrical stimulation parameters can be tuned to achieve therapeutically beneficial neural activity that can differ between cell types (*i*.*e*. stable activity in one population and reduced activity in another)(26). Therefore, it is possible that modulating stimulation waveform may be able to increase the therapeutic window of DBS through differential activation of subpopulations (Fig. 4-5, Fig. S2) and frequency-dependent progression of population activity (Fig. 6E-F). However, the connectivity and cellular composition influences the population responses during DBS(83) and these factors differ considerably between deep brain structures and the cortex(84-86). Therefore, it will be important to experimentally determine how waveform asymmetry modulates activation of neural subtypes in the cortex and deeper brain structures(87) and how their respective activity changes over timescales relevant to symptom relief(88).

### Limitations

This study was conducted in layer II/III of sensory cortex in anesthetized mice with an upward facing Michigan-style electrode. Because ketamine anesthesia influences baseline neural activity and synaptic transmission(89-91), it is possible that spatial and temporal patterns of stimulation-induced calcium activity may differ in an awake preparation. Additionally, although waveforms were presented in a random order, it is possible that preferential activation is influenced by anesthesia and plasticity(92). The electrode geometry influences the distribution of charge and electric field(63, 93), therefore the spatial distribution of direct and indirect activation may be different with conical electrodes like the Utah array, which are used clinically(4, 6). Additionally, the distribution of neuron subtypes with different geometries changes throughout the cortex(55, 94, 95) and the response to ICMS can be layer dependent(96). Therefore, although some of the electrodes on a Utah array may be in layer II/III(97), the progression of population activity is likely to differ for electrodes in different layers(96, 98), or brain regions(75). The mouse model used in this study expresses the calcium indicator only in a subset of excitatory neurons(31); however, stimulation is likely to modulate the activity of inhibitory neurons as well as glial cells(10, 99, 100) which may contribute to the observed patterns of activity. Future studies should further investigate how stimulation waveform and frequency modulate the activity of these other cell types in an awake preparation.

### Conclusions

Intracortical microstimulation has the potential to restore sensory perception in neural prosthetics. Our data support the hypothesis that direct activation of neural elements occurs within 20-60 μm of the electrode and demonstrates that waveform asymmetry may modulate the direct activation of different neural populations. Importantly, this waveform-and frequency-dependent activation of different neural populations differentially modulated the temporal progression of population activity, which highlights the complexity of ICMS parameter space and suggests further investigation into how diverse neural populations respond. Therefore, stimulation waveform represents a valuable parameter in ICMS that merits further investigation for neural prosthetics.

## Conflict of Interest

The authors declare no competing interests. KAL is a scientific board member and has stock interests in NeuroOne Medical Inc., a company developing next generation epilepsy monitoring devices. KAL is also paid member of the scientific advisory board of Cala Health, Blackfynn, Abbott and Battelle. KAL also is a paid consultant for Galvani and Boston Scientific. KAL is a consultant to and co-founder of Neuronoff Inc. None of these associations are directly relevant to the work presented in this manuscript.

## Author contributions

Conceptualization: KCS, JRE, KAL, TDYK. Methodology: KCS, JRE, KAL, TDYK. Investigation: KCS. Formal analysis: KCS. Visualization: KCS. Writing – Original Draft: KCS, TDYK. Writing – Review & Editing: KCS, JRE, KAL, TDYK. Funding Acquisition: TDYK.

## Acknowledgements

We want to thank Steven Wellman, Keying Chen, Fan Li, and Christopher Hughes for critical review of the manuscript. This work was supported by: NIH R01NS094396, R01NS105691, R01NS115707, NSF CAREER 1943906, ARCS Foundation Inc., Pittsburgh Chapter, Gookin Family Foundation Award and Stover Family Foundation Award

## Research Data

The data that support the findings of this study are available from the corresponding author upon reasonable request.

## Supplementary Figures

**Figure S1.**
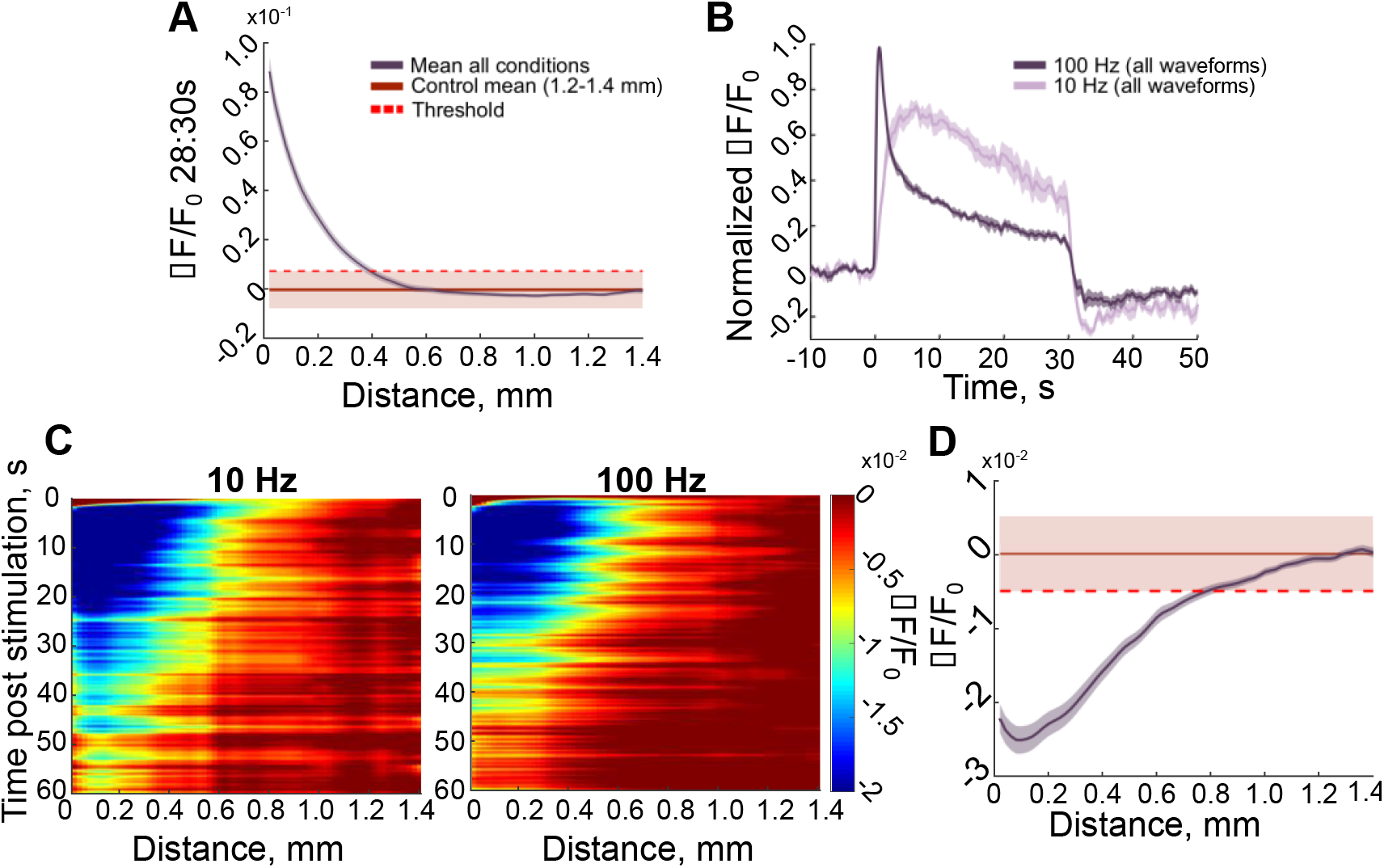
Stimulation induced activity is concentrated near the electrode. **A**. Stimulation induced neural activity (purple) at the end of stimulation is focused within the first 400 μm around the electrode. The threshold is the mean plus one standard deviation over all groups (all 8 waveforms and both 10 Hz and 100 Hz) within 1200-1400 μm from the electrode (red). **B**. There is substantial reduction in normalized neural activity soon after the onset of 100 Hz-stimulation (0.5-1.5s; dark purple) followed by a slower reduction in activity (1.5-30s). This slower reduction in activity (1.5-30s) is comparable to the reduction in activity during 10 Hz-stimulation (light purple). Data were averaged over the first 400 μm **C**. The end of stimulation is followed by a long-lasting and wide-spread period of depressed neural activity (dark blue; 3-20 s post stimulation). Calcium activity reduces below baseline levels beginning ∼3s following the end of stimulation and lasts for ∼17s. **D**. The long-lasting depression of activity (purple) extends farther than the area of activity that is substantially elevated during stimulation (680 μm vs 400 μm). Activity was averaged during 3-20s post stimulation. Control activity (red) is the average activity within a control region 1200-1400μm from the electrode during 3-20s post stimulation for all groups. Shaded regions for stimulation activity (purple) represents standard error, and the shaded region for the control activity (red) is standard deviation to represent the threshold calculation.

**Figure S2.**
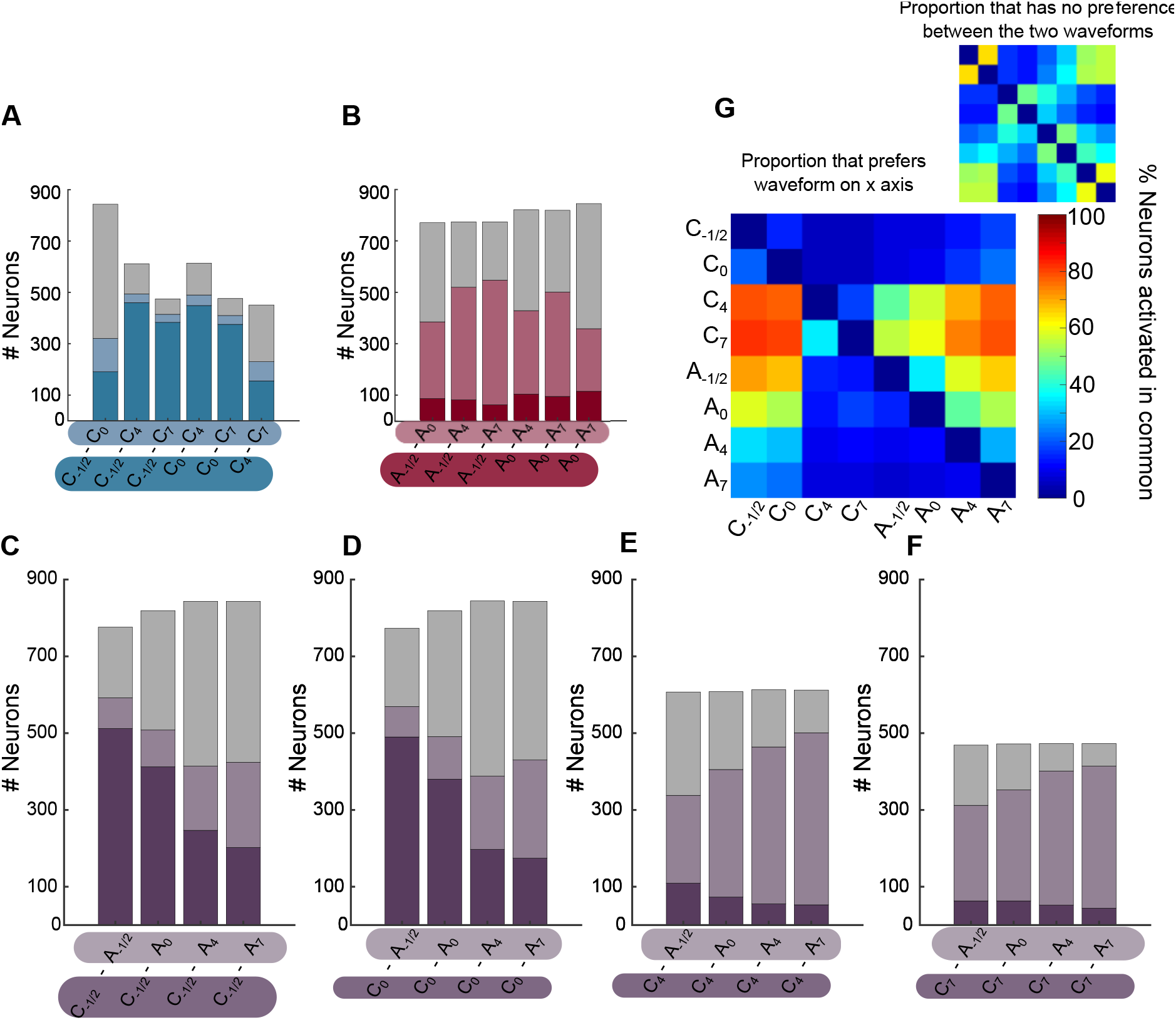
Certain neural populations are preferentially activated by specific waveforms over others. A-F. Representation of the proportion of all neurons activated in common between two specific stimulation waveforms being compared that are preferentially activated by one waveform over the other (shaded colors), or do not display preference between the two (Grey). **G**. The proportions for all comparisons in A-F represented as a heatmap to highlight trends between waveforms. Neurons are generally preferentially activated by symmetric cathodal-first pulses compared to asymmetric cathodal-first (A) or asymmetric anodal-first compared to symmetric anodal-first pulses (B). Neurons seem to have similar activation for cathodal-first stimulation pulses with longer return phase or symmetric cathodal-first stimulation pulses compared to asymmetric anodal-first stimulation with long activation phase (C-D). Similarly, neurons are often preferentially activated by anodal-first stimulation pulses compared to asymmetric cathodal-first stimulation pulses (E-F). Although neurons generally display this preferential activation, these results highlight that waveform is modulating the population activation differently than simple amplitude changes.

